# MolMeDB: Molecules on Membranes Database

**DOI:** 10.1101/472167

**Authors:** Jakub Juračka, Martin Šrejber, Michaela Melíková, Václav Bazgier, Karel Berka

**Affiliations:** Regional Centre of Advanced Technologies and Materials, Department of Physical Chemistry, Faculty of Science, Palacký University Olomouc, tř. 17, listopadu 12, 771 46 Olomouc, Czech Republic

## Abstract

Biological membranes act as barriers or reservoirs for many compounds within the human body. As such, they play an important role in pharmacokinetics and pharmacodynamics of drugs and other molecular species. Until now, most membrane/drug interactions have been inferred from simple partitioning between octanol and water phases. However, the observed variability in membrane composition and among compounds themselves stretches beyond such simplification as there are multiple drug-membrane interactions. Numerous experimental and theoretical approaches are used to determine the molecule-membrane interactions with variable accuracy, but there is no open resource for their critical comparison. For this reason, we have built Molecules on Membranes Database (MolMeDB), which gathers data about over 3600 compound-membrane interactions including partitioning, penetration, and positioning. The data have been collected from scientific articles published in peer-reviewed journals and complemented by inhouse calculations from high-throughput COSMOmic approach to set up a baseline for further comparison. The data in MolMeDB are fully searchable and browsable by means of name, SMILES, membrane, method, or dataset and we offer the collected data openly for further reuse and we are open to further additions. MolMeDB can be a powerful tool that could help researchers better understand the role of membranes and to compare individual approaches used for the study of molecule/membrane interactions.

**Database URL:** http://molmedb.upol.cz.

## Introduction

Biological membranes consist of complex lipid and protein mixtures that play a crucial role in molecular transport into/out of cells. Apart from passive or active permeation, molecules can also accumulate in the membranes at specific functional positions or they can disrupt the membrane altogether. All those molecule-membrane interactions are important for the actions of individual molecules in the organism and their pharmacokinetics.

And yet, most chemical databases use octanol/water partition coefficient (logP) as the only measure of small molecule interactions with lipid membranes, but the membrane compositions of individual cells and organelles can widely vary as it is being currently unraveled by findings from lipidomics (1). The membrane protein structural databases provide additional information not only about the position and the topology of the membrane proteins but also about the membrane type localization (e.g., OPM (2), PDBTM (3), MemProtDB (4), TPML (5), or EncoMPASS (6)); however, the data about various molecule/membrane interactions are scattered among different sources. For example, DrugBank (7) covers logP and information about membrane transporters and carriers for many drug molecules, but it does not provide a measure for the assessment of penetration nor does it involve partitioning through individual membranes. Permeability of compounds through skin membranes is either present in EDETOX database (8) or scattered throughout literature, e.g., sources cited in supplemental information of ref. (9). Similarly, the recently established PerMM database (10) covers only cellular permeability together with permeability prediction using an implicit membrane model with rigid compounds. Finally, molecular dynamics simulations are often used for predictions of membrane partitioning (11) or permeability even on a large scale (12,13). However, current theoretical predictions of molecule/membrane interactions vary by method as well as in comparison with data from experiments, lacking community benchmark comparison between individual methods.

To fill this gap, we have developed Molecules on Membranes Database (MolMeDB) as an open and up-to-date online manually curated depository of molecule/membrane interactions. MolMeDB contains over 3600 interactions described in the literature or obtained by our COSMOmic-based high-throughput calculations (14). In addition to listing the individual molecule/membrane interactions, we provide a simple tool for comparison of interactions between multiple methods and/or membranes. Using this information, it is possible to analyze the membrane behavior of the selected subsets of molecules. Examples of these analyses are provided as case studies to better illustrate efficient ways to extract useful knowledge from the MolMeDB database.

## Materials and Methods

### Data Collection

To collect datasets of molecule-membrane interactions, a manual inspection of articles (15–23) and already existing databases (e.g., PerMM database (10)) with the focus on expressions like ‘membrane partition coefficient’, ‘membrane permeability’ or ‘permeability coefficient’ was performed (Figure 1). Primarily, we focused on high-throughput experimental setups like Black Lipid Membrane (BLM) (24), Parallel Artificial Membrane Permeability Assay (PAMPA) (17,25), Caco-2 permeability assay (22), liposomal fluorescence assay (20), *n*-hexane passive dosing (26), and polydimethylsiloxane (PDMS) based permeabilities (19,27) that provide partition coefficients of compounds on a variety of natural and artificial membranes. Moreover, we have also collected resources from a broad variety of computational methods, e.g., molecular dynamics-based Umbrella Sampling (US) approach, COSMO-RS theory-based COSMOmic calculations, or implicit solvent-based PerMM model. Those methods also differ in the level of approximation or force fields used to predict the compounds properties. This diversity within individual methods provides additional verification, especially in comparison to experimental data.

**Figure 1.**
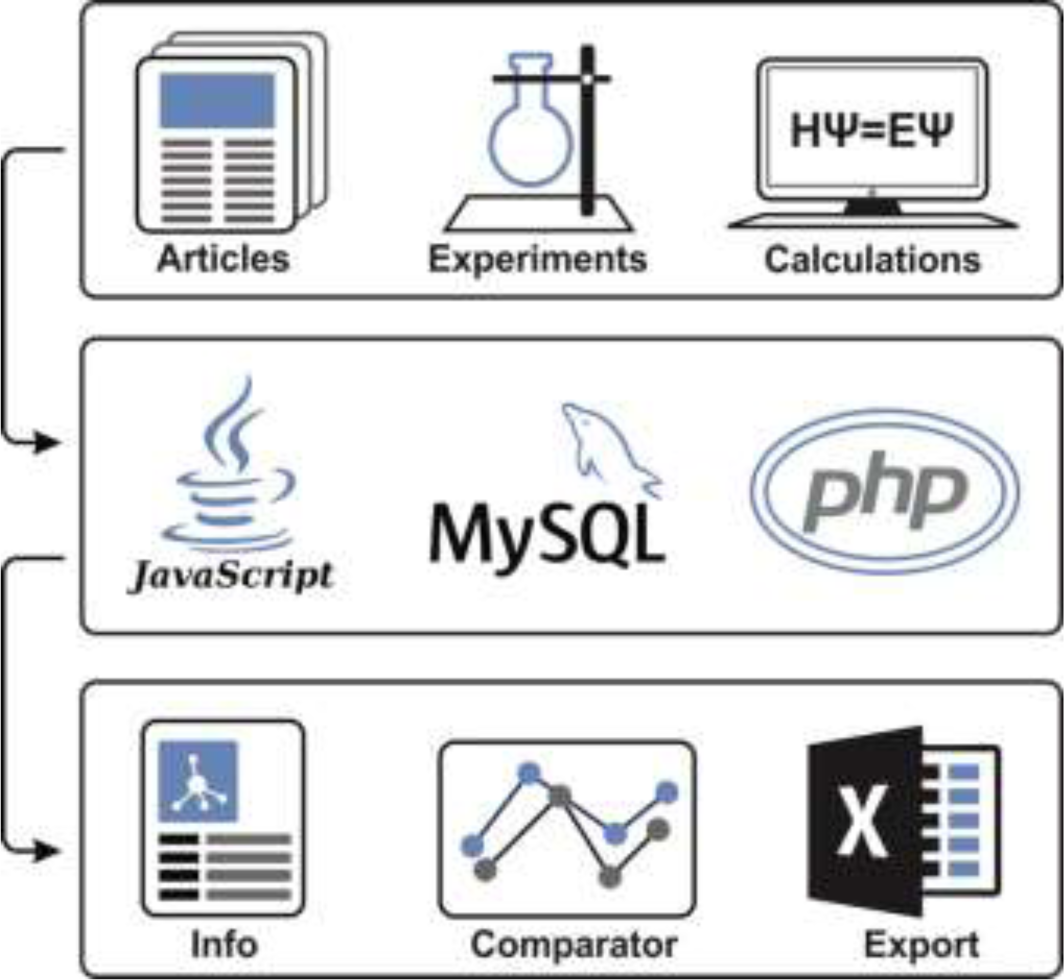
Illustrative scheme of MolMeDB workflow. Input data collected from experimental/theoretic studies are curated and introduced into the MySQL database with a web interface in HTML5/CSS + PHP7. Data for individual molecule/membrane interactions are visualized either as data tables or in interactive JavaScript graphs, and they can be directly compared and downloaded.

### In-House COSMOmic Calculations

Apart from the already published data, we have added our original dataset of XY compounds on various membranes mimicking either cell-like membranes (DMPC or DOPC bilayers) or skin-like membranes (ceramide NS or *stratum corneum* mixture bilayers consisting of an equimolar mixture of ceramide NS:cholesterol:lignoceric acid).

Neutral conformers of compounds were generated from SMILES with the LigPrep and MacroModel modules (Small-Molecule Drug Discovery Suite 2015-4, Schrödinger, LLC, New York, NY, 2016, https://www.schrodinger.com). Individual conformers of each compound were generated using the OPLS_2005 force field (28) in vacuum. Mixed MCMM/LMC2 conformational searches were performed to enable low-mode conformation searching with Monte Carlo structure selection. Maximum ten conformers were selected for further analysis if they were within 5 kcal/mol of the lowest energy conformer and, to reduce the number of similar conformers, had an atom-positional RMSD of at least 2 Å relative to all other selected conformers. Each selected conformer was subjected to a series of DFT/B-P/cc-TZVP vacuum and COSMO optimizations using Turbomole 6.3 (Turbomole V6.3 2011, http://www.turbomole.com) within the cuby4 framework (29). After each optimization step, single point energy calculations at the DFT/B-P/cc-TZVPD level with a fine grid (30) were performed to obtain COSMO files for each conformer. The structures of COSMO .mic files describing bilayers were then obtained from fitting COSMO files of individual lipids to the bilayer structures obtained from free 200 ns+ long molecular simulations from refs for DMPC (11), for DOPC and ceramide NS (31) and for *stratum corneum* mixture bilayers (32). For each conformer/lipid sets we calculated free energy profiles using COSMOmic 15 (14) or COSMOmic/COSMOperm 18 to obtain averaged free energy profiles. From those, information about membrane partitioning, permeability and affinity, central energy barrier, and the position of drug at its energetical minima was extracted.

#### Database Architecture

MolMeDB webpage is built with the combination of HTML5/CSS and PHP7 layouts running on Apache server. The database runs on MySQL (Figure 2). The AJAX search engine allows search over names of compounds, datasets, or SMILES. 2D structures are generated from SMILES using CDK Depict (33). 3D structure visualization is provided by LiteMol (34) over MOL files generated via RDkit (RDKit: Cheminformatics and Machine Learning Software. 2018, http://www.rdkit.org), or downloaded from PubChem (35), or DrugBank (7) databases, or uploaded by the user. DrugBank is used also for interconnection links to other databases. Free energy profiles of individual molecule/method/membrane sets, where available, are visualized using Chart.js JavaScript application (https://chartjs.org). PMF profiles data are stored as equidistant values spaced 1 Å for possible comparison between individual PMF profiles and they are interpolated from the uploaded data by Neville’s algorithm of iterated interpolation.

**Figure 2.**
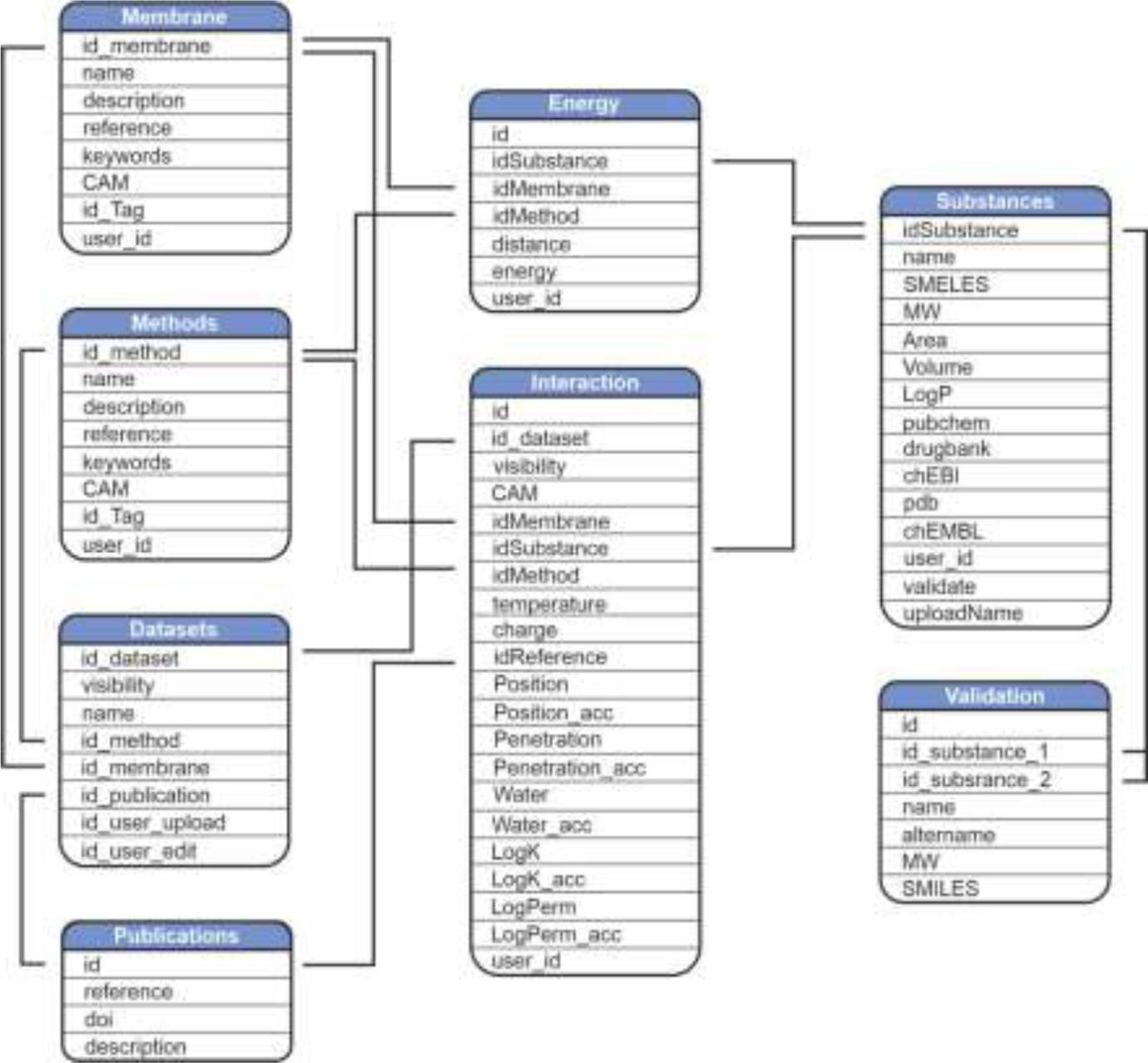
Scheme of MolMeDB database.

### Results and Discussion

#### User Interface and Database Usage

To provide a user-friendly interface for the molecule-membrane interaction data, we developed a web version of MolMeDB database, freely accessible at http://molmedb.upol.cz. Interaction data can be accessed via browse or search functions. ‘Browse’ section (Figure 3A) allows the user to scroll through the list of available compounds, membranes, and methods used to describe the molecule-membrane interactions. ‘Search’ section (Figure 3B) enables to search for a desired compound by its name or SMILES notation and to look up compounds measured/computed by individual methods and membranes. Dataset search allows the user to browse within a list of publications by title or authors’ names. All data can be added into the Comparator tool (described below). MolMeDB web also includes ‘Documentation’ section (Figure 3C) explaining the methodology, giving several examples and a tutorial for using the database. Finally, ‘Statistics’ section (Figure 3D) keeps track of the number of entries and of interactions in subsets of individual methods or membranes.

**Figure 3.**
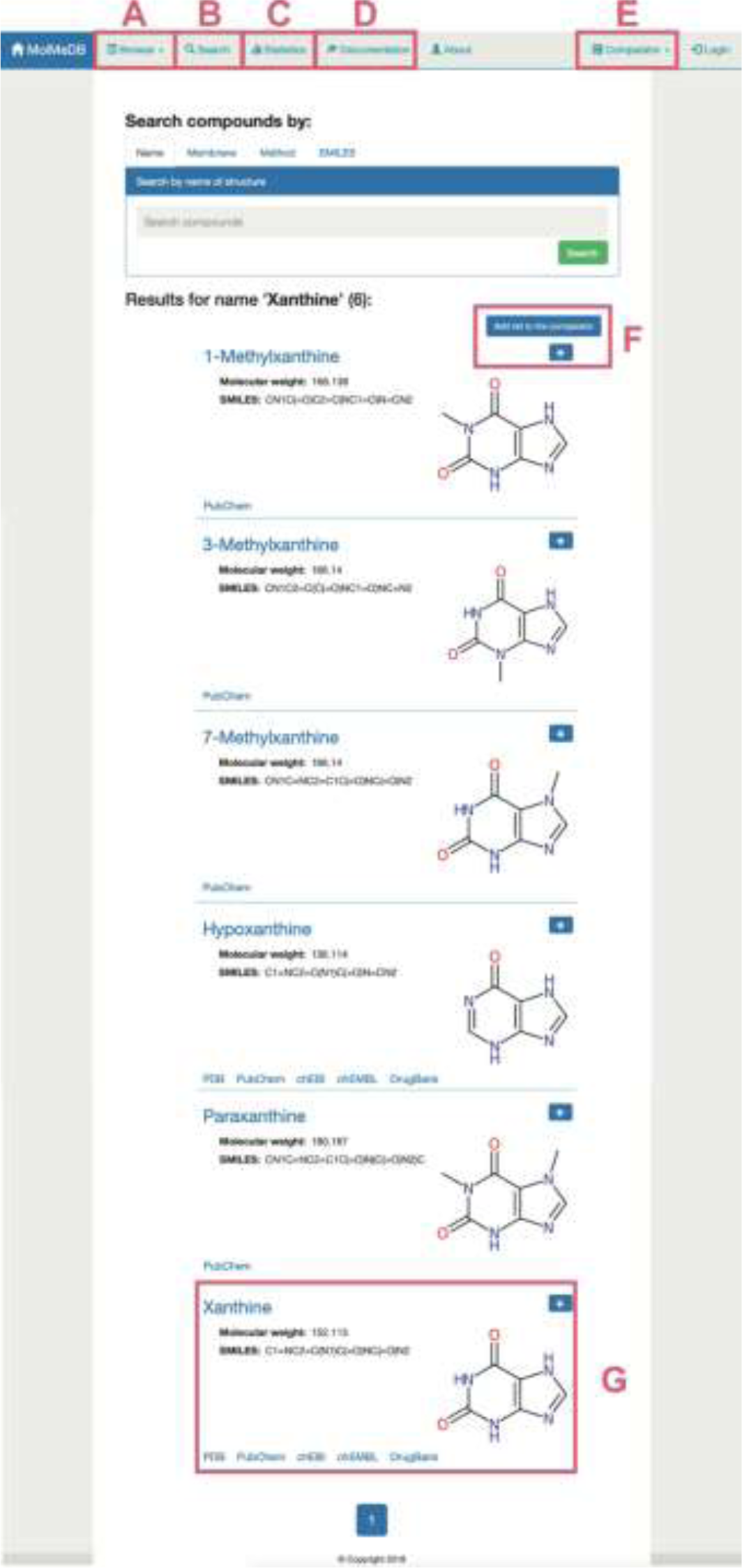
Layout of a page showing the toolbar of Browse/Search (A, B) utility along with menu items for Statistics (C), Documentation (D) and Comparator tool (E). Example of search utility for “xanthine” molecule. Compounds with corresponding pattern of name are selected and displayed along with 2D structure (G). Target molecules can be directly added into molecule Comparator (F) by clicking on “+” sign.

The user can use the ‘Search’ section (Figure 3A) to list all compounds matching the entry name. As example of “xanthine” entry, 6 compounds were listed partially matching the given expression (Figure 3). Among the listed entries, the user gets a quick overview of the compounds properties, 2D structure, and links to other chemical databases containing information about the molecule (e.g., Protein Data Bank (36), PubChem (35), ChEBI (37), ChEMBL (38), DrugBank (7)). Desired molecules can be added into Comparator tool (see below). After selecting a compound, in our case “xanthine”, a purine-based molecule which serves as a parent compound for caffeine and its derivatives, the page is divided into three sections (Figure 4):

1. General info (Figure 4A)—provides a description of compound properties like molecular weight along with links to other databases via the molecule’s identifiers. The first section also shows a 2D image generated from SMILES via CDK Depict and a 3D structure generated by RDKit or downloaded from PubChem or DrugBank databases visualized with LiteMol.
2. Interactions table (Figure 4B)—displays an interactive table with molecule-membrane interactions such as membrane/water partitioning (logK_m_), permeability coefficient (logP_erm_), free energy barrier in the membrane center (ΔG_pen_), affinity towards the membrane measured by the energetical minimum (ΔG_wat_), or the position of the interaction minimum for the molecule on membrane (Z_min_) available for a combination of membranes and methods with variation where available. Charge of compound (Q) and temperature are specified for each molecule-membrane interaction. The user can then switch among individual methods to compare the measured/calculated properties. Source references are also listed in Publication field. The desired data can be directly downloaded as a .csv table.
3. Free energy profile graph (Figure 4C)—demonstrates the course of the free energy profile along the membrane normal with energetical barriers and minima between membrane center (0 nm) and water environment (3.5 nm). For an individual method, the user can switch among available membranes and directly visualize given energetical profiles.

**Figure 4.**
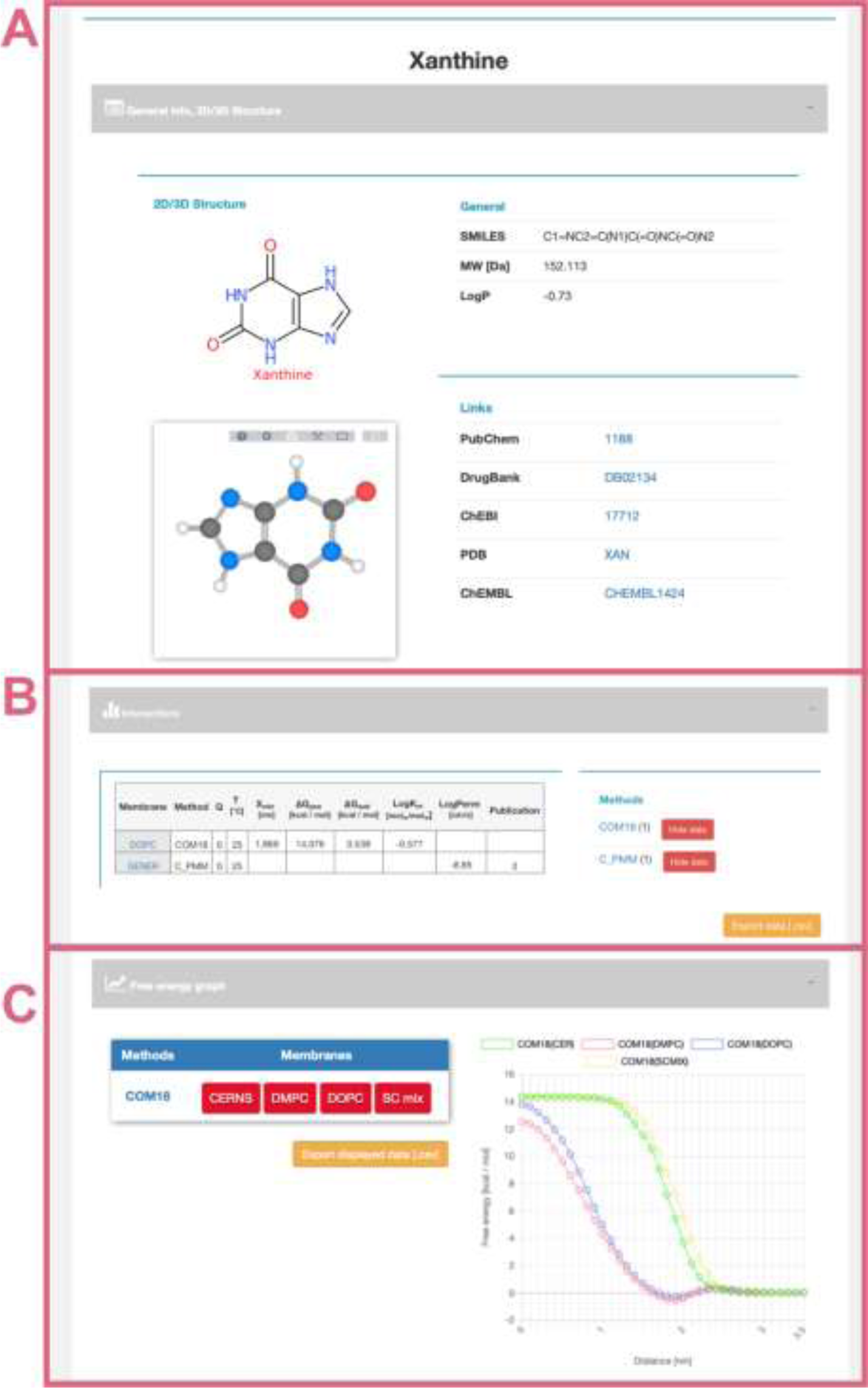
Interface of selected “xanthine” molecule divided into separate panels for general information (A) about the molecule with its 2D and 3D structure and links to other databases; interactive table showing available data about molecule-membrane interactions (B) on position, partitioning, energy barrier, and permeability coefficient for a given pair of method/membrane and charge; and interactive graph with available free energy profiles (C).

#### Use Cases - Comparator Tool

Although interaction data for individual compounds are valuable as such, their comparison allows the use of the data for research within multiple scientific fields. For this purpose, we have embedded the Comparator tool which allows to gather molecule-membrane interaction data for multiple compounds from one or more methods and to compare them in order to visualize patterns within the data or to assess the validity of predictive methods.

##### Caffeine and its Metabolites

Caffeine is a purine-based molecule, which is metabolized in a set of multiple-step reactions into a series of chemically modified compounds. In the first step, caffeine is metabolized to the following metabolites: theobromine (by enzymes CYP1A2, CYP2E1), theophylline (CYP1A2, CYP2E1), 1,3,7-trimethyluric acid (XO), 6-amino-5(*N*-formylmethylamino)-1,3-dimethyluracil (CYP1A2), and paraxanthine (CYP1A2) (Figure 5) (39,40).

**Figure 5.**
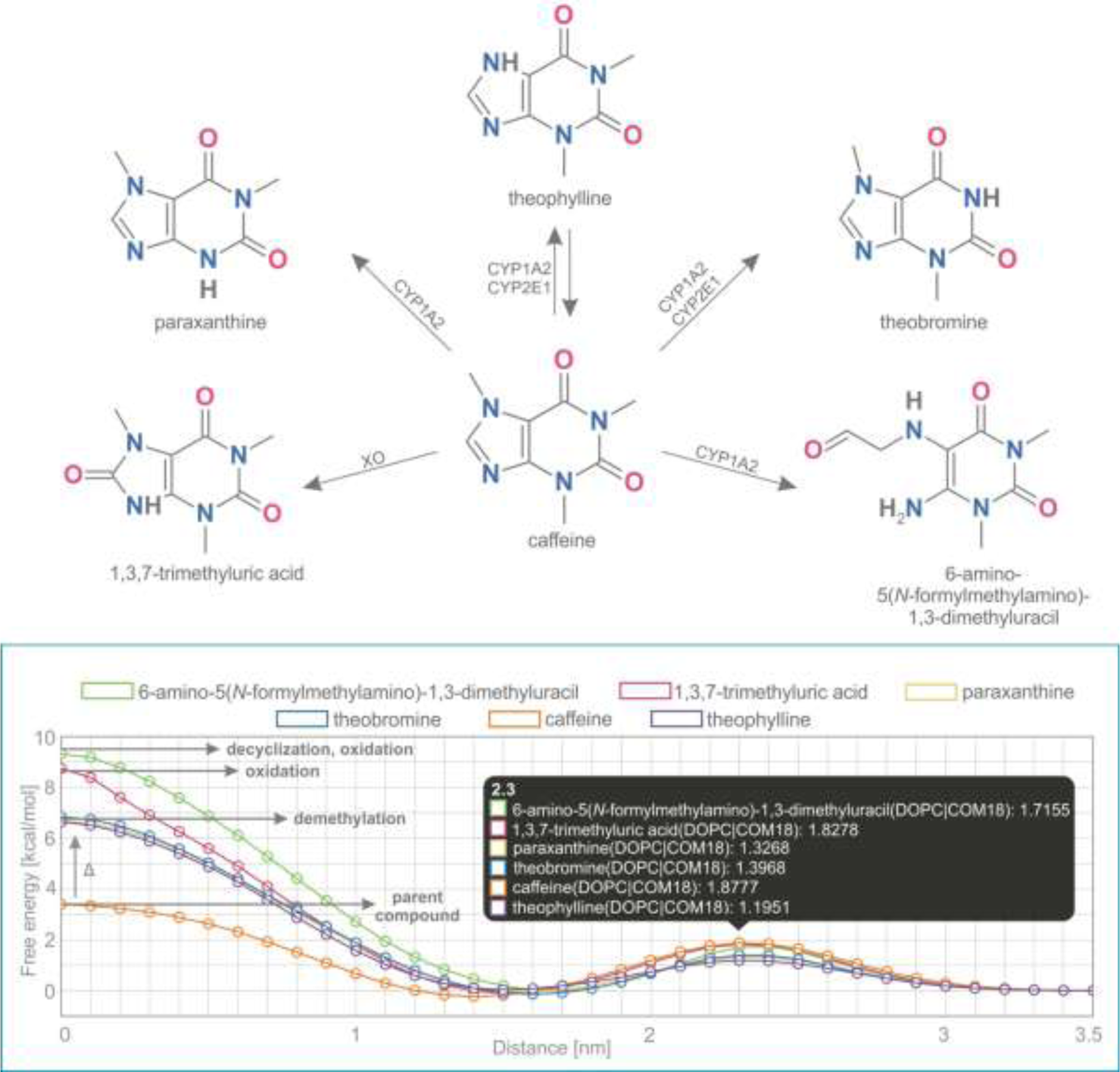
Comparison of free energy profiles of caffeine and its metabolites showing an increase of penetration barrier according to the type of metabolizing reaction and to the nature of chemical modification.

Free energy profiles on DOPC membrane for this set of caffeine derivatives were calculated using COSMOmic 18. The nature of the chemical modification of the parent molecule caused different interactions of the metabolites with the membrane. Caffeine derivatives are metabolized in three distinct types of reactions: demethylation (theobromine, theophylline, paraxanthine), oxidation (6-amino-5(*N*-formylmethylamino)-1,3-dimethyluracil, 1,3,7-trimethyluric acid), and decyclization (6-amino-5(*N*-formylmethylamino)-1,3-dimethyluracil). Here we show that individual types of caffeine metabolites exhibit distinguishable changes in free energy profiles (Figure 5). The lowest energetic barrier (ΔG_pen_) is shown for caffeine as the parent compound, followed by all demethylated metabolites with an identical increase of penetration barrier by approximately 3.4 kcal/mol. Oxidized and oxidized/decyclized products experienced an even greater increase of penetration barrier compared with the original caffeine molecule by 5.4 and 5.9 kcal/mol, respectively.

Overall, all products of caffeine metabolism show a hindered passage through the membrane core as evidenced by the increase in the penetration barrier energy (ΔG_pen_). On the other hand, the affinity of all molecules towards the membrane (ΔG_wat_) remained almost the same, which is in concord with their very similar logP values. Finally, all metabolites shifted their energetic minima (Z_min_) toward the membrane/water interface by 2 Å.

#### Comparison of Methods

The Comparator tool also allows a comparison of multiple methods/membranes with each other over selected compounds. Such type of comparison can be used to evaluate different theoretical approaches (e.g., PerMM, COM18) versus experimental data (e.g., BLM).

In this example, we show a comparison of permeability coefficient datasets obtained from theoretical PerMM and COSMOmic/COSMOperm 18 predictions and experimental BLM method for 101 neutral/unionized compounds on DOPC/generic phosphatidylcholine membrane. The user can reach the whole dataset for an individual Method/Membrane via the Search tool and add it directly into the Comparator tool. Upon choosing the desired combination of multiple Method/Membrane options and charge of molecules, the data can be plotted in an interactive window (Figure 6). The individual permeability coefficient for a particular compound can be visualized by hovering the cursor over the given datapoint (here an examples for Citric acid and Hydrofluoric acid). Linear regression fit was used here to determine the level of correlation between the two methods, obtaining the coefficient of determination (R^2^) of of 0.76 and 0.77 for COM18/BLM and PerMM/BLM respectively, while PerMM data show almost identical slope to the experimental data.

**Figure 6.**
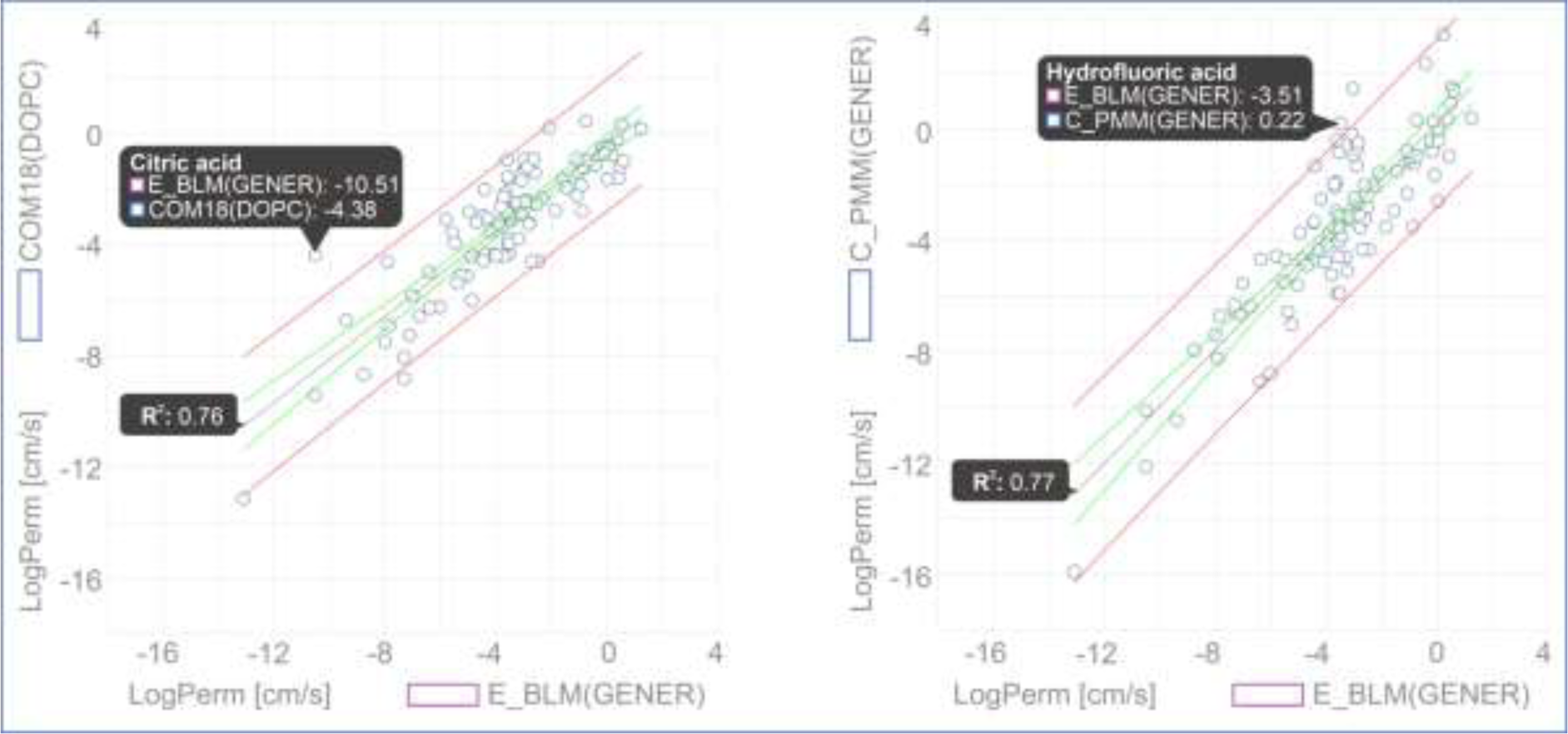
Comparison of permeability coefficients obtained from experimental BLM method and theoretical PerMM model. The figure was manipulated externally from downloaded data to add linear regression line shown in grey with confidence interval shown in red and prediction interval, in green. Confidence interval was set to 95%.

## Conclusion and Future Work

MolMeDB is a unique, manually curated database on interactions of compounds with membranes. To date it contains more than 1200 compounds and 3600 molecule-membrane interactions obtained both theoretically and experimentally. MolMeDB stores multiple descriptors of molecule/membrane interactions and provides the tools for searching and browsing these data and their comparison. MolMeDB can prove to be a valuable resource for many research groups to benchmark the key data on molecule-membrane interactions, which are important in the fields of pharmacology, toxicology, and molecular simulations. In the future, we plan to add further datasets and implement also the involvement of transporters and carriers. More complex statistical analysis within the selected datasets and further FAIRification of the data is anticipated in following versions of MolMeDB. We believe that MolMeDB can be a useful starting point, which can facilitate future studies devoted to a deeper understanding of biological roles of molecules on membranes and that it will attract the biological membrane community to establish a common ground for sharing open data in this field.

## Availability and requirements

MolMeDB database is freely available at http://molmedb.upol.cz. The visualization of 3D structures with the LiteMol molecular viewer requires the browser to have WebGL enabled.

